# Computational Design of a New Aflatoxin B1 Aptamer *in lieu* of SELEX Technique

**DOI:** 10.1101/2022.11.12.513184

**Authors:** Mohamad Yasser Ahmad Ghazy

**Affiliations:** Research and Development Department, Mounir Armanious Research Center, Eva Pharma, Giza, Egypt

**Keywords:** APTAMERS, APTASENSORS, COMPUTATIONAL BIOLOGY, MD, MOLECULAR MODELLING, SELEX, MAKING APTAMERS WITHOUT SELEX

## Abstract

Mycotoxins are extremely dangerous, and their detection in our environment, food and feed is becoming increasingly important. Biosensors are being implemented heavily in mycotoxin detection along with other significant applications. Aptamers have numerous beneficial advantages as biorecognition molecules and are being used as the biorecognition part of biosensors (Aptasensors). The development of aptamers does not require inducing immune response against the target, but the SELEX method is used. The SELEX method is laborious, time consuming and can be expensive at times. Various efforts were done to replace that method with a computational alternative to reduce the effort, time and money needed to develop and design aptamers. One of the most significant efforts to achieve that was the MAWS algorithm. We used the MAWS algorithm to develop a new aptamer against aflatoxin B1, the most dangerous mycotoxin. The MAWS algorithm failed to function properly, and molecular modelling and molecular docking was used alternatively to achieve the same goal. A new pipeline for predicting ssDNA aptamers was proposed, a new aptamer against aflatoxin B1 was obtained and recommendations for further future research directions were given.

## I. Introduction

Mycotoxicosis is a toxic effect exerted on animals, humans and other living organisms by mycotoxins, low-molecular weight organic chemical compounds that are widely present in the food chain. The toxic effect of mycotoxins can be both acute and chronic. Their ingestion can cause serious problems, such as kidney and liver damage, immunosuppression, nervous system diseases, carcinogenicity and mutagenicity **(Moez *et al*., 2020)**. Mycotoxins are produced by filamentous fungi, such as *Penicillium, Fusarium*, and *Aspergillus* species, which strengthen the fact that these toxins will be present whenever the conditions are favorable for the growth of these fungi **(Adeyeye, 2016)**. The presence of mycotoxins is mostly noticed in soil, animal products, animal feedstock and most importantly, human food. Beside their toxic effect on humans and animals’ health, they possess a toxic effect on the international trade **(Tola and Kebede, 2016)**. This is due to the reduction of yield and quality of products contaminated with these toxins, such as milk, coffee, meat, cereals, dried fruits.

Accordingly, mycotoxins gained international attention and the MPLSs (maximum permissible limits) and LOD (limit of detection of analytical techniques) were set by numerous organizations, such as USFDA, EU, FAO and WHO, in an effort to ensure safety and sustainability. Due to the serious effects of mycotoxins, the LOD is very low and low PPB’s (Part Per Billion) Concentrations are used. Simultaneous multitoxin detection and determination can be done with methods that possess superior analytical capability and has very low LOD. Examples of these conventional methods are MS (mass spectrometry), LC-MS/MS (Liquid chromatography-tandem-MS), GC-MS (Gas chromatography-MS), UV absorption detectors, TLC (Thin layer chromatography), HPLC with FLD (High performance liquid chromatography with fluorescence detector), ELISA (Enzyme-Linked Immunosorbent Assays) and FT-NIR (Fourier Transform near-infrared). However, the main problems with Mycotoxin analysis methods are summarized in **(Figure 1.1) (Malhotra, 2014; Vidal et al., 2013)**.

**Figure (1.1).**
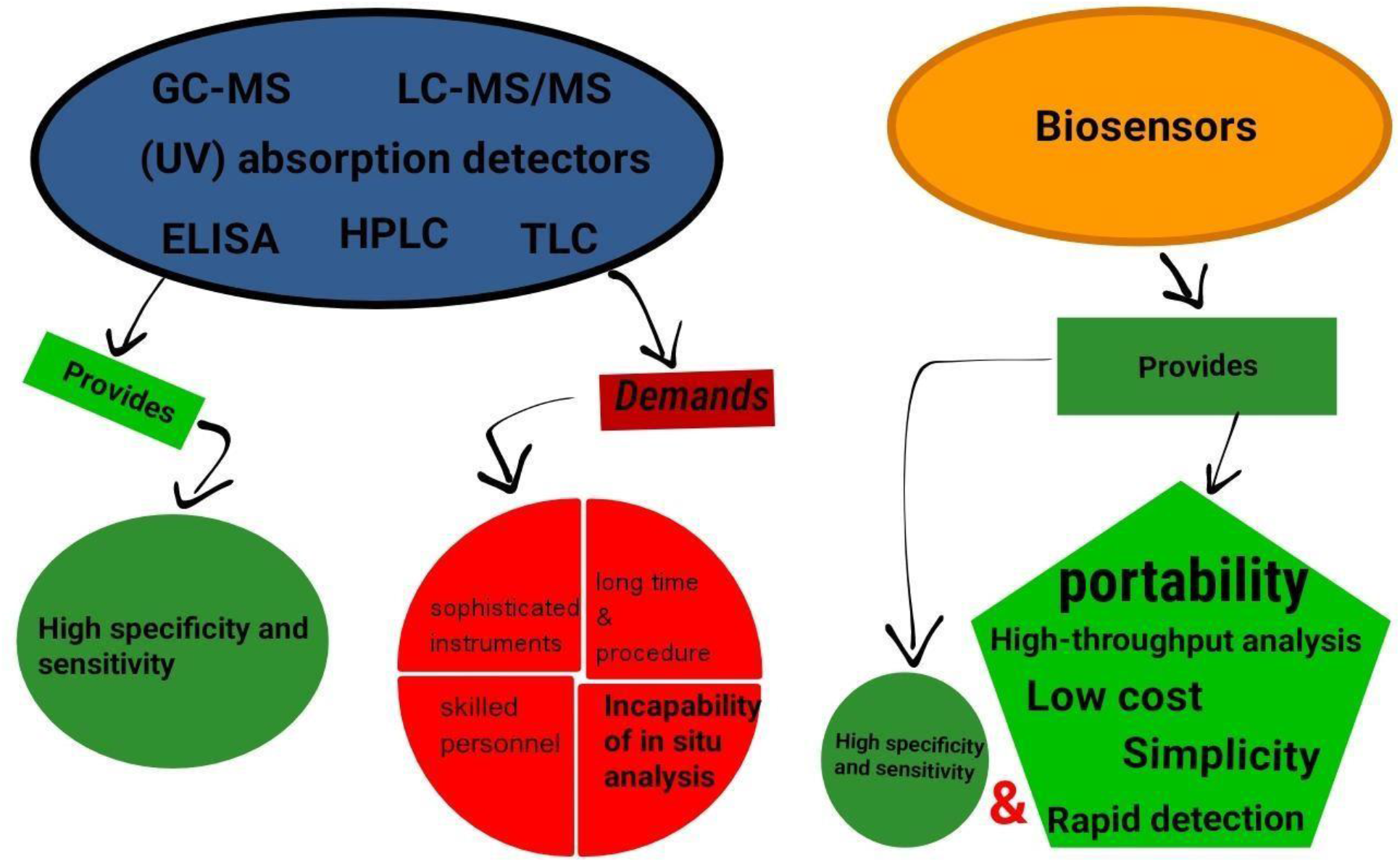
Biosensors vs conventional techniques for mycotoxins analysis and detection.

On the other hand, biosensors emerged as an innovative as well as promising alternative tool to the conventional techniques for mycotoxins detection. Beside their capability to fulfill the LOD set by legalizations, biosensors possess superior advantages to the conventional methods, such as their portability, rapid detection, high-throughput analysis with noticeably low cost, and their remarkable simplicity, whether in their manufacturing or their operating, without compromising with the required specificity and sensitivity **(Figure 1.1) (Anene et al., 2016)**.

Generally, biosensors consist of three main parts, biorecognition component, immobilization matrix and a transducer to convert the reaction product into a measurable signal **(Figure 1.2)**. The most crucial part of any biosensor is the biorecognition one, various types of molecules can be used to detect mycotoxins such as antibodies, enzymes, DNA, RNA or synthesized single stranded oligonucleotides (DNA or RNA) known as Aptamers. Antibodies possess high susceptibility to heat, and irreversible denaturation and their high molecular weight make their usage in biosensors somewhat problematic. Consequently, it was highly desirable to find an alternative for proteins in general as biorecognition molecules, i.e., aptamers. Aptamers have the ability to bind to a large range of targets with high stability, affinity, and specificity. They became more favorable due to their reproducibility, small molecular weight which allows high density detection, high stability which allows them to be reused repeatedly with no loss in their binding ability, high binding affinity, high specificity and the easiness of their modification that enables them to have a high binding affinity for any given molecule **(Fernández et al., 2017)**.

**Figure (1.2).**
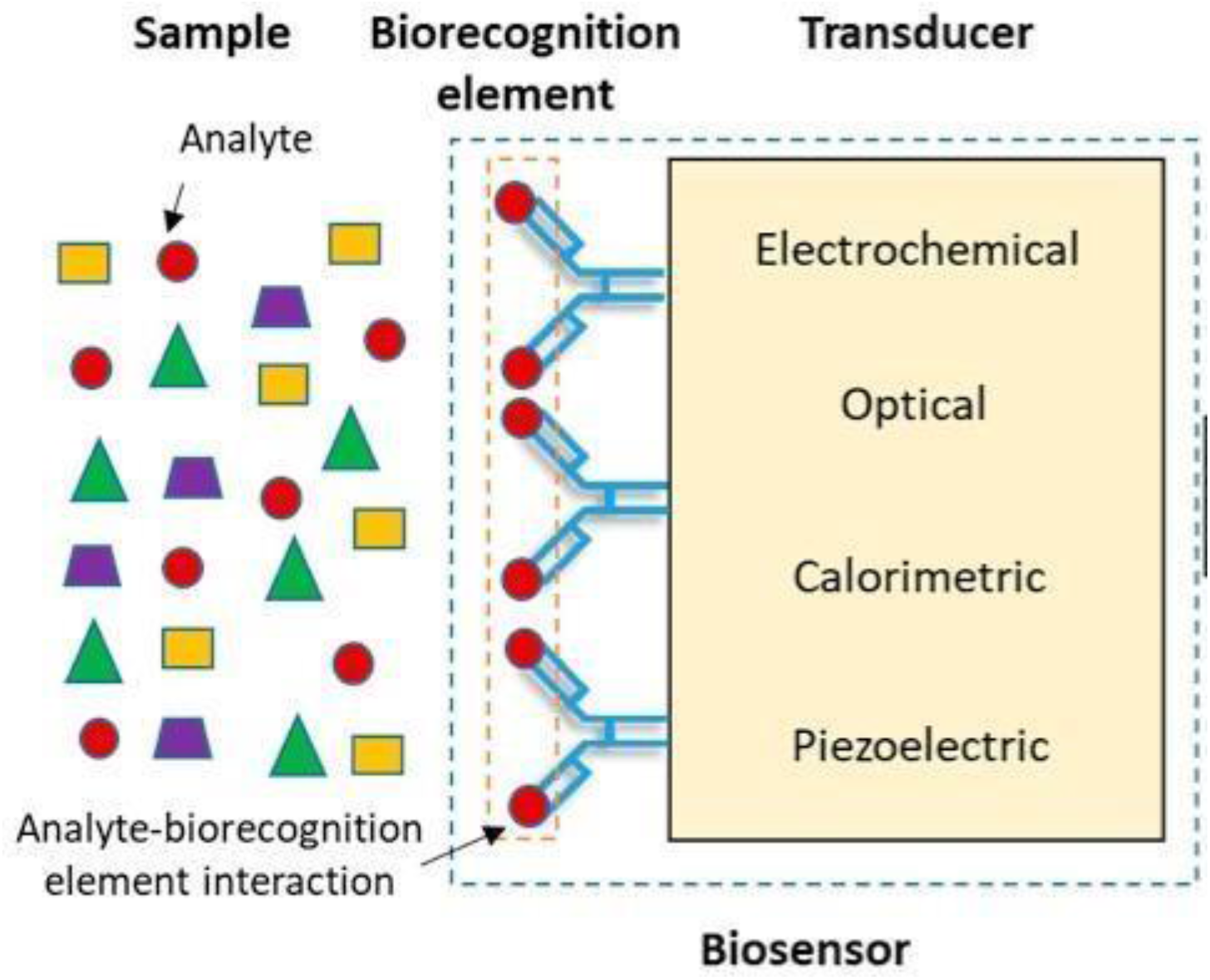
Simple representative diagram of Biosensors general composition **(Contreras-Naranjo and Aguilar, 2019)**.

Most aptamers are obtained through SELEX, systematic evolution of ligands by exponential enrichment, which is a process that requires months to be completed and requires a highly amount of handling. The technique begins by producing convenient oligonucleotide sequences library which generally consists of 10^13^ to 10^15^ random DNA or RNA sequences. The generated library is incubated with the immobilized target molecule and the unbound strands are washed away. On the other hand, the bound sequences are separated and then amplified by PCR. Single strands of the amplified sequences are obtained and seeded for the next round. The previous steps are repeated 10 to 12 times. Besides these rounds, negative SELEX and counter SELEX rounds, where the aptamers are incubated with other targets that are either similar to the primer target or not, are added for ensuring and enhancing the specificity of the selected aptamer. Obtaining a specific aptamer is never guaranteed after all of these rounds nor is obtaining an aptamer in the first place **(Younis *et al*., 2020)**.

In the age of technology and computational biology, performing that tedious process to obtain an aptamer is impractical. We aim to obtain an aptamer specific for Aflatoxin B1, the most dangerous mycotoxin, through a computational approach without resorting to performing SELEX.

## II. Review of Literature

In the recent years, biosensors, which are analytical devices that convert a biological response into an electrical signal, have evolved. Typically, biosensors must be stable, i.e., physical parameters such as temperature and pH should not tamper with it, and it also must be highly specific, portable, and reusable. The Construction of biosensors, its transducing devices, materials, and immobilization techniques, requires multidisciplinary research in engineering, biology, and chemistry. The materials used in biosensors are classified into three classes based on their processes:

1 biocatalytic group (e.g., Enzymes)
2 bioaffinity group (e.g., antibodies and nucleic acids)
3 Whole cells (e.g., microorganisms).

Biosensors applications is vast and of great importance. In this section we will be discussing some of these applications and their composition, particularly, the different types of biorecognition molecules. The Aptasensors and the development of aptamers will be focused on along with the mention of the importance and urgency of detecting mycotoxins in foodstuff, specially, aflatoxin B1.

### 1. Biosensors

Biosensors are devices used to detect the presence or concentration of a biological analyte, such as a biomolecule, a biological structure, or a microorganism. Biosensors consist of three parts: a component that recognizes the analyte and produces a signal, a signal transducer, and a reader device and **(Moez *et al*., 2020)**.

#### 1.1 Biosensors applications

Biosensors have been applied in many fields namely, medical field, food industry etc., and they provide great advantages over the traditional methods with high sensitivity and stability **(Mehrotra, 2016)**.

##### 1.1.1 Medical field

The applications of biosensors are expanding swiftly in the medical field. For an excellent example, Glucose biosensors. Glucose biosensors are extensively used in clinical applications for the monitoring and diagnosis of *diabetes mellitus* **(Scognamiglio *et al*., 2010)**. Due to the ease of operation of biosensors and the fact they do not require experienced labor to function properly, 85% of the gigantic world market is accounted for blood-glucose biosensors that can be used at home **(Mehrotra, 2016)**. Biosensors are being used prevalently in the diagnoses of transmittable diseases. It also has cardiac-related applications **(Lee *et al*., 2012)**. Biosensors are capable of biochemical recognition with high selectivity for the desired biomarker. The numerous other biosensors applications include neurochemical detection by diamond microneedle electrodes, microfluidic impedance assay for controlling endothelin-induced cardiac hypertrophy, quantitative measurement of cardiac markers in undiluted serum, immunosensor array for clinical immunophenotyping of acute leukemia and biochip for an accurate, yet rapid, detection of multiple cancer markers **(Mehrotra, 2016)**.

##### 1.1.2. Biodefense biosensing applications

Biosensors are used in the rapid and accurate detection of BWAs (Biowarfare agents), e.g., namely, bacteria (vegetative and spores), toxins and viruses, in order to avoid or deal with biological attacks. The main purpose of such biosensors is to identify organisms posing threat in virtually real time with extremely high sensitively and selectively. One given example is the use of biosensors in the detection of HPVs, the human papilloma virus HPV (double stranded DNA virus) which is related to invasive cervical cancer. This probe directly detects HPV genome with no need to the polymerase chain reaction amplification process **(Mehrotra, 2016)**.

##### 1.1.3 Metabolic engineering

There is an urging need for the development of the microbial synthesis of chemicals. Researchers view metabolic engineering as the empowering technology for a maintainable bioeconomy **(Woolston *et al*., 2013)**. They have also proposed that a significant portion of pharmaceuticals, fuels, chemicals, and commodity will be produced from renewable feedstocks by using microorganisms rather than count on petroleum refining or extractions from plants. To achieve such goal, an efficient screening methods is required to select the individuals carrying the desired trait. However, the methods used for that purpose had limited throughput, e.g., spectroscopy-based enzymatic assay analytics. To dodge this hindrance, genetically programmed biosensors that facilitate *in vivo* observing of cellular metabolism were created to present potential for high-throughput selection and screening using cell survival and fluorescence-activated cell sorting (FACS), respectively **(Peroza *et al*., 2015)**.

##### 1.1.4. Plant biology

Biosensors can be utilized to identify missing components pertinent to metabolism, regulation, or transport of the analyte. Roger Tsien’s lab was first to develop protein prototype sensors to measure caspase activity and control levels of calcium in live cells **(Jones *et al*., 2013)**.

##### 1.1.5 food processing, monitoring, food authenticity, quality, and safety

A tough impasse in food processing industry is of quality and safety, Traditional techniques are exhausting, time consuming and expensive. Hence the need for an alternative with consistent and objective measurement for food monitoring and authentication of food products, in a cost-effective manner, is highly existing for the food industry. Thus, development of biosensors in response to the demand for an inexpensive, simple yet highly selective and real-time techniques is seemingly propitious **(Scognamiglio *et al*., 2014)**.

1 monitoring of ageing of beer (enzymatic biosensors) **(Ghasemi-Varnamkhasti *et al*., 2012**)
2 Detection of pathogens in food (potentiometric alternating biosensing systems) **(Arora *et al*., 2011**)
3 organophosphate pesticides detection in milk (automated flow-based Enzymatic biosensors) **(Mishra *et al*., 2012)**
4 fermentation processes Biosensors can be utilized to monitor the presence of products, biomass, enzyme, antibody, or by-products of the process to indirectly measure the process conditions. Nowadays, several Kinds of commercial biosensors are accessible that can detect biochemical parameters (glucose, lactate, lysine, ethanol etc.) and capable of online monitoring of fermentation processes. Moreover, they are widely used in China, occupying about 90% of its market **(Yan *et al*., 2014)**.
6 Biosensing technology for sustainable food safety Biosensors are used to detect pesticides in wine and orange juice **(Suprun *et al*., 2005)**. Moreover, Arsenic can be measured with the help of bacteria-based bioassays **(Diesel *et al*., 2009)**.
7 Mycotoxins detection

As the content of mycotoxins is particularly low in food, the development of probes to detect AFB1 in foods with high sensitivity and selectivity is an urgent social need for the evaluation of food quality **(Tola and Kebede, 2016)**. Numerous techniques have been developed to monitor AFB1. Nevertheless, most of them require labor-consuming, and sophisticated instruments, which have limited their application **(Anene et al., 2016)**.

#### 1.2. Biosensors components and their types

The bioreceptor and the transducer are argued to be the most important components of any biosensor. The type of the physicochemical signal produced through the reaction between the bioreceptor, and the analyte varies, and the form of the transducer used varies accordingly. Whether, electrochemical, mass, thermal, or optical, the transduction element can come in various forms **(Wondergem *et al*., 2017)**. The important components of a biosensor are (1) a bioreceptor (e.g., enzymes, antibody, microorganism, or cells); (2) a transducer of the physicochemical signal, and (3) a signal processor to interpret the information that has been converted.

As mentioned, bioreceptor can be classified as:

##### 1.2.1. Enzymes

Enzymes are able to retain their catalytic activity within certain parameters within a limited range. These parameters are mainly, chemical agents, fluid forces, temperature, and pH. However, they are well suited for potentiometric and amperometric biosensors giving to their high turnover numbers. Although enzymes are widely used in biosensing systems, they have a high-cost factor in terms of purification and production as they must be produced from biological sources (**Monosik *et al*., 2012)**.

##### 1.2.2. Antibodies

Antibodies, also Known as immunoglobulins (Ig), are a class of globular proteins (∼150 Kda) which bind to specific antigenic structures forming an antibody-antigen complex, and they have widespread applications in clinical diagnostics and biotechnology. Their basic structure comprises of four polypeptide chains arranged in a Y configuration. The two binding sites on the arms of the antibody can be extremely variable, allowing millions of different antibodies with slightly varying paratopes to exist. This lends antibodies to be widely applied in immunoassays for molecule detection across cells, large proteins, and even amino acids (**Monosik *et al*., 2012)**.

##### 1.2.3. Cells and Microorganisms

An interesting and growing area of research which uses this principle is that of synthetic biology, where a commonly used microbial chassis (e.g., E. coli, S. cerevisiae) is transfected with a gene that is turned on in the presence of a specific substrate. This has rapid sensing applications in healthcare monitoring of viruses and toxins **(Lim *et al*., 2015)**.

Microbial biosensors are classified, based on the sensing method, into two main groups:

A. electrochemical microbial biosensors
B. optical microbial biosensors.

Owing to their massive production through cell culturing, microbes (such as algae, bacteria, or yeast) are used in biosensor systems. Enzymatic biosensors are more sensitive than microbial biosensors and the overall detection period are shorter than that of microbial biosensors, however microbial biosensors are well suited to environmental analysis due to their ability to detect the global effect of some toxin not just a specific compound or toxin **(Lim *et al*., 2015)**.

##### 1.2.4. Nucleic Acids

Nucleic acid-based sensing systems are more sensitive than antibody-based detection methods as they provide gene-based specificity, without utilizing amplification steps to attain detection sensitivity to the required levels. Aptamer generation has been achieved for a wide variety of targets including small molecules **(Jhaveri and Ellington, 2002)**, proteins **(Bardoczy and Meszaros, 2006)**, viruses **(Balogh *et al*., 2010)**, and whole cells **(Sefah *et al*., 2010)**. Unlike antibodies, aptamers can also be generated for targets that are toxic as well as for targets that do not elicit an immune response *in vivo* **(Jayasena, 1999)**. Additionally, aptamer sequences can be modified with reporter molecules; this allows for labeling at judiciously chosen nucleotide positions to minimize any effect on the functionality of the aptamer **(Beaucage and Lyer, 1993)**.

## 2. Aptasensors

DNA and RNA aptamers are single-stranded oligonucleotides which fold into secondary and tertiary structures forming specific binding pockets for low or macro molecular compounds of different types, including cells, cell surface proteins, bacteria, and viruses, under certain conditions (ionic strength, pH, temperature). Their specificity is equivalent to that of antibodies and in some cases, even higher. The RNA selection method with improved enzymatic activity to cleave DNA has been identified by **(Robertson and Joyce, 1990)**. That work also introduced the term “aptamer” (from the Latin aptus, meaning” to fit” and Greek meros, meaning “the part”).

### 2.1. SELEX

Aptamers are identified based on a combinatorial approach as previously discussed, that approach is SELEX. DNA binding with ligands, such as proteins or other compounds, consists of a variety. The stability of complexes is defined by the apparent constant of dissociation, Kd. Kd usually ranges in the pM-nM range for aptamer-protein complexes, which is equivalent to that of antibody-antigen complexes. For instance, platelet-derived growth factor subunit B DNA aptamer Kd is 100 pM, and vascular endothelial growth factor (VEGF) RNA aptamer Kd is about 50 pM. However, for small molecule-aptamer complexes, a higher Kd (nM-mM) is normal **(Bardoczy, 2002) (Jhaveri, 2006)**. Unbound molecules of DNA/RNA are eluted from the column, while bound aptamers are isolated and then amplified by PCR from the complex. This cycle is replicated several times (around 6-10) and the result is that the sequence of DNA or RNA is obtained with a high affinity to the target ligand. However, the SELEX does not provide the aptamers with the highest affinity for the target substratum in all situations, considering the high variety of sequences. Post-SELEX modification can therefore assist in improving binding properties. Currently, several SELEX improvements have been recorded. Among these, negative or counter SELEX increases the affinity of aptamers and removes those in a dynamic setting of non-specific bindings to other molecules **(Younis *et al*., 2020)**.

### 2.2. RNA Aptamers vs DNA Aptamers

DNA and RNA aptamers’ affinity to their ligands is comparable. RNA aptamers have more variable structures, however. On the other hand, in complex biological liquids, RNA aptamers are less stable in contrast to DNA aptamers. In ideal solvent conditions and with sufficient chemical modification, the aptamers are very stable and can shape a number of structures. Typically, guanine-rich aptamers form guanine tetrads stabilized by the so-called Hogsten bonds. The tetrads can be organized parallel by their plane and form guanine quadruplexes (G-quadruplexes) **(Macaya *et al*., 1993)**. The charge density of the quadruplex is typically twice higher in comparison with those of linear nucleic acids. This is advantageous in electrostatic interactions with positive charged binding sites at the molecule, for example, thrombin. G-quadruplexes are typical for the aptamers that specifically bind thrombin, VEGF, or ochratoxin A **(Macaya *et al*., 1993)**.

### 2.3. Advantages of using Aptamers

The distinctive value of aptamers is their outstanding structural reproducibility. If the aptamer sequence is chosen, using well-established oligonucleotide chemistry, it can be replicated at any time with high accuracy. Aptamers are fairly versatile and can be modified by different ligands, which enhance their stability in complex biological liquids and allow them to be immobilized on different surfaces. Biotin, thiols, amino or carboxyl modified linkers are among convenient Kinds of modifications **(Fernández et al., 2017)**. Furthermore, aptamers can be updated by groups of reporters that significantly enhance their properties as receptors in biosensors. Since the discovery of the aptamer, the comprehensive application of aptamers as receptors began in biosensors **(Younis *et al*., 2020)**.

### 2.4. Aptamers for small molecules

Despite this trend towards larger targets, there are many compelling reasons for pursuing the identification of new small molecule-binding aptamers. Small molecules play Key roles in many biological processes due to their ability to diffuse across cell membranes **(Cho and Juliano, 1996)**. These targets may be harmful and carcinogens as toxins, or beneficial, such as drugs or nutrients. In cells, small molecules serve as cell signaling molecules, pigments, or as part of defense mechanisms **(Ashour and M. Wink, 2011)**.

Giving the upmost importance of the detection of Aflatoxin B1 (AFB1), one of the most commonly existing mycotoxins in contaminated food, and the great applicability and the highly advantageous properties of aptamers, Aptasensors are considered to be an emerging strategy for the quantification of aflatoxin B1 with high selectivity and sensitivity **(Roemer *et al*., 2012)**.

## 3. Computational applications in Aptamer selection

Aptamers are mostly identified through an iterative process called systematic evolution of ligands by exponential enrichment (SELEX), which is a cyclic process that involves multiple rounds of selection and amplification **(Tuerk and Gold, 1990)**. The entire process is tedious, time consuming and often fails to enrich high affinity aptamers **(Osborne and Ellington, 1997)**. Additionally, the requirement of fixed priming sites in the sequences of a library imposes a length criterion on random region, thereby restrict the diversity of the synthesized aptamer library. Even the lengthy aptamers, usually ≥40 nt long require prior trimming and shortening for their efficient scale-up production **(Ahirwar *et al*., 2016)**.

### 3.1. Binding mechanism determination

The recent progress in these selection methods allows now to fulfill the increasing demand in selective receptors for targets of interest (cells, viruses, proteins, and small molecules) in different areas ranging from therapeutics **(Fang and Tan, 2010)** to diagnosis **(Cho *et al*., 2009)** and sensing **(Hayat and Marty, 2014) (Zhou *et al*., 2014)**. However, in spite of the large number of aptamers selected against a wide range of molecules, only a few efforts have been made to characterize the binding mechanism between aptamers and their target, such studies being performed only for a restricted number of popular molecules of interest **(Challier *et al*., 2016)**. still, mechanistic characterization of the binding process through the determination of structural, thermodynamic, and Kinetic properties are obviously a major prerequisite in order to rationally design further applications. For instance, many of the aptamer-based analytical detection strategies involve either the labeling of the aptamer oligonucleotide (i.e., by a molecular probe, a protein, a nanoparticle, etc.) or the anchoring of the 3’ or 5’ extremity of the aptamer to a surface (aptasensor). Such labeling requires at least Key information concerning the binding mechanism in order to optimize its impact on the recognition process **(Challier *et al*., 2016)**.

Single-stranded oligonucleotides often possess readily ascertainable structures that may predefine a target binding site, and most of the aptamers studied so far adopt such a prestructuration due either to high content in canonical base pairing, theophylline **(Stojanovic *et al*., 2001)** and cocaine **(Zimmermann *et al*., 1997)**, or to the formation of a G-quadruplex structure (thrombin) **(Macaya *et al*., 1993)**. For those systems, the correlation between NMR, nuclear magnetic resonance, and molecular dynamics has proven its efficiency for the aptamer/target complex mapping **(Buck *et al*., 2007; Huizenga and Szostak, 1995)**. Conversely, in the critical case for which there is no trivial pre-structuration of the aptamer, no predefined binding site and no accessible structural information on the final aptamer/target complex, elucidation of the binding mechanism is even more difficult to achieve **(Challier *et al*., 2016)**.

### 3.2. Computational SELEX simulation

During nucleic acid modeling and *in silico* design, a full set of *in silico* methods can be applied, such as docking, molecular dynamics (MD), and statistical analysis. The typical modeling workflow starts with structure prediction. Then, docking of target and aptamer is performed. Next, MD simulations are performed, which allows for an evaluation of the stability of aptamer/ligand complexes and determination of the binding energies with higher accuracy. Then, aptamer/ligand interactions are analyzed, and mutations of studied aptamers made. Subsequently, the whole procedure of molecular modeling can be reiterated. Thus, the interactions between aptamers and their ligands are complex and difficult to understand using only experimental approaches. Docking and MD are irreplaceable when aptamers are studied *in silico* **(Buglak *et al*., 2020)**.

#### 3.2.1 ZDOCK-ZRANK

Computational simulations of aptamer-protein interactions were carried out by using ZDOCK and ZRANK functions in Discovery Studio 3.5 starting from the available information of aptamers generated through the systematic evolution of ligands by exponential enrichment (SELEX) in the literature. From the best of three aptamers on the basis of ZRANK scores, 189 sequences with two-point mutations were created and simulated with Ang2. Then, they used a surface plasmon resonance (SPR) biosensor to test 3 mutant sequences of high ZRANK scores along with a high and a low affinity binding sequence as reported in the literature. They found a selected RNA aptamer has a higher binding affinity and SPR response than a reported sequence with the highest affinity. This is the first study of *in silico* selection of aptamers against Ang2 by using the ZRANK scoring function, which should help to increase the efficiency of selecting aptamers with high target-binding ability **(Hu *et al*., 2015)**.

The ZDOCK algorithm in Discovery Studio uses the fast Fourier transform (FFT) correlation techniques and searches all possible binding positions of the two proteins. The original scoring function of ZDOCK is a geometrical measure according to the degree of shape complementarity between the two docking proteins. For the software package DS 3.5, the ZDOCK program incorporates the pairwise shape complementarity scoring function for identifying docked conformations and scores hits based on atomic contact energies. In the past study, the degree of shape complementarity (SC) was used to quantify the interaction between two interacting surfaces **(Lawrence and Colman, 1993)**, and the thrombin-aptamer complex had better SC than most antibody/antigen interactions **(long *et al*., 2008)**. Based on the calculation of shape complementarity, we used the ZDOCK algorithm to study the aptamer-Ang2 interaction. In addition, the ZRANK scoring function was also applied to each simulation in this study that reranked initial-stage ZDOCK predictions with detailed electrostatics, van der Waals, and desolvation energy terms. In our previous research, we adopted the ZDOCK method to simulate the interactions between the thrombin and aptamers, and simulation results were consistent with experimental results from literatures **(Kumar *et al*., 2013)**.

#### 3.2.2 Dubbed Mutation Monte Carlo (MMC)

MMC is a computational method that has been published to enable mapping of such sequence preferences **(Eslami-Mossallam *et al*., 2016)**. Dubbed Mutation Monte Carlo (MMC), the method utilizes standard Monte Carlo simulations to sample the Boltzmann distribution associated to a modeled DNA system such as the nucleosome and adds as a novel feature Monte Carlo moves that mutate the DNA sequence. Given a suitable model of the system of interest, this technique allows an understanding of the sequence preferences of the system from a theoretical point of view. The MMC method shares many similarities with the experimental SELEX method. It samples DNA sequences based on their affinity to the target. Doing so at constant finite temperature delivers probability distributions for, e.g., dinucleotides **(Tompitak *et al*., 2017)**, and by performing simulated annealing it searches for the sequence with the strongest affinity **(Tompitak *et al*., 2016)**, much as Lowary and Widom, **(1998)** attempted, leading to the 601 sequence. However, there is also a major difference between the *in silico* method and the experimental protocols. The MMC simulation is performed at a particular temperature, which determines how stringently it selects for low-energy states and hence for high-affinity sequences. This temperature is necessarily shared by both the configurational moves that simulate the thermal fluctuations of the system and the mutations. In a SELEX experiment, however, the selection pressure is determined by, among other factors, the number of rounds of selection performed, and the strength of selection on the sequences is decoupled from the temperature at which the experiment is performed. Despite the similarities, this means that a MMC simulation cannot be directly taken as an *in-silico* SELEX experiment **(wondergem *et al*., 2017)**.

#### 3.2.3 Step by Step Mutation based on MD simulation

A computational method to design a new aptamer with higher binding affinity to a special target in comparison with the experimentally available aptamers. The method is called step by step mutation based on MD simulation, which includes some steps. First, MD simulation is performed for the SELEX-introduced (native) aptamer in the presence of the target. Afterwards, conformational factor (Pi) is calculated for the simulated system, which obtains the affinity of the aptamer residues to the target. A nucleotide exchange is done for the residue with the least Pi parameter to the nucleotide with the highest Pi value that results in a mutant aptamer. MD simulation is performed for the target-mutant complex, and Pi values are calculated again. The nucleotide exchange is performed similarly, and the designing process is proceeded repeatedly that results in a mutant with the improved specificity to the target. The aptamer affinity to the target is also determined in each step through calculating the binding Gibbs energy (ΔGBind) as a reliable parameter. The introduced strategy is utilized efficiently to design a mutant aptamer with improved specificity toward sulfadimethoxine (SDM) antibiotic as a case study. The great difference in the ΔGBind values about 579.856 KdJ mol−1 highlights that the M5 mutant possesses the improved specificity toward SDM in comparison with the native aptamer. Besides, the selectivity of the M5 aptamer toward SDM is examined among some conventional interfering compounds by using MD simulation that confirms the applicability of the designed aptamer for further experimental studies **(Khoshbin and Housaindokht, 2020)**.

#### 3.2.4 Designing 3D structure of ssDNA aptamers

An approach to design the stem-loop/hairpin for the three-dimensional structure of DNA aptamers through serial applications of computational prediction methods by comparing the simulated structures with the experimental data deposited in PDB Data bank, followed by MD simulations was suggested by **(Sabri et al, 2020)**. The result shows minimal structural differences were observed between the designed and the original NMR aptamers, and the stem-loop conformational structures were also retained during the MD thus suggesting the proposed aptamers designing methods are able to synthesize a high-quality molecular structure of hairpin aptamers, comparable to the NMR structures.

#### 3.2.5 In silico Docking-assisted SELEX

Successfully selected a ssDNA aptamer, Z100, against Zearalenone (ZEN) after 8 rounds of SELEX was selected successfully by (**Zhang *et al*., 2018)**. Further, *in silico* docking assisted study was performed to investigate the binding mechanism between ZEN and Z100 using AutoDock Vina open-source program ver. 1.1.2. The binding free energy was approximated at −9.6 Kdcal/mol. **Jokar *et al*., (2016)** carried out molecular docking integrated experiments using the ssDNA aptamer and acetamiprid. The AutoDock results of acetamiprid-aptamer showed 6.812 ± 0.533 Kdcal/mol of binding energy. As for the binding site, the docking results also revealed that SS2-55 and SS4-54 loops as the active sites of aptamer and confirmed the formation of the aptamer-Acetamiprid complex. To summarize, molecular docking provides a deeper understanding of aptamer-target interaction, which is useful for the optimization and rational design of DNA aptamers **(Tereshko *et al*., 2003)**. Various reliable molecular docking simulation programs are available such as GRAMM **(Bruno *et al*., 2011)**, Hex **(Hu *et al*., 2019)**, ZDOCK **(Yarizadeh *et al*., 2019)**, HADDOCK **(Khan *et al*., 2019)**, PatchDock **(Bruno, 2019)**, NPDocK **(KaKoti and Goswami, 2016)**, AutoDockTools **(Mousivand, 2020)**, AutoDock **(Jokar *et al*., 2016; Zavyalova *et al*., 2020)**. Most of the docking programs are internet-based servers or easily downloadable programs useful to the scientific community **(Navien *et al*., 2021)**.

#### 3.2.6 Microarray-assisted method

Microarray aptamer analysis was combined with *in silico* secondary structure prediction. *In silico* studies applied three conditions to the aptamers: (1) presence of a predicted secondary DNA structure producing one hairpin loop without a ligand, (2) hairpin loop with a length from 3 to 7 bases, and (3) stem length from 6 to 9 bases. As a result, a novel patulin aptamer was optimized **(Tomita *et al*, 2016)**.

#### 3.2.7 SELEXION

In 1991, SELEX was first simulated using a program named SELEXION (Systematic Evolution of Ligands by Exponential Enrichment with Integrated Optimization by Non-linear Analysis). SELEXION was first used to reconstruct bacteriophage T4 DNA polymerase gp43 SELEX experiments **(Irvine *et al*., 1991)**.

#### 3.2.8 AptaSim

A program to simulate the aptamer selection process called “AptaSim” was coded by **(Hoinka *et al*., 2015)**. AptaSim aimed at realistically recreating the selection process during SELEX with the intention of investigating the effect of error-prone PCR on aptamer selection **(kinghorn *et al*., 2017)**.

#### 3.2.9 Improving affinity of aptamers

Bioinformatic approaches have been used to improve the affinity of aptamers. As highlighted earlier, due to low selection pressure classical SELEX is unlikely to resolve the very best aptamer sequences. Therefore, each individual aptamer generated using a bioinformatics approach must be individually assayed for binding affinity, which can be labor-intensive and time-consuming. DNA microarrays consist of many features or spots on a glass slide, each feature containing many copies of a unique DNA sequence. This high-throughput technology allows for simultaneous assay of many aptamer sequences via incubation with fluorescent target **(kinghorn *et al*., 2017)**. There are several software packages and databases customized for aptamer scientists to analyze the large amount of HTS data based on different strategies **(kinghorn *et al*., 2017)**.

The use of a patterned library in SELEX was able to select specific binders for all three molecules with affinity at nanomolar levels better than those selected from random libraries (streptavidin: KD = 105 nM, IgE: KD = 26 nM, VEGF: KD = 45 nM). These results showed that the use of a patterned library could increase the proportion of active aptamer, speed of selection, and affinity of the resultant aptamers **(Ruff *et al*., 2010)**.

In conclusion, Bioinformatic approaches have been used to improve both aptamers and their selection. We outlined a broad range of aptamer bioinformatics techniques including simulation of aptamer selection, aptamer selection by molecular dynamics, patterning of libraries, identification of lead aptamers from HTS data, and *in silico* aptamer optimization.

### 3.3. Computational alternatives to SELEX

Generally, the computational prediction approaches of aptamer have been proposed to carry out in two main categories: interaction-based prediction and structure-based predictions **(Emami *et al*., 2020)**.

Several different approaches have been developed to simplify the whole selection process. Most notable of these are *in silico* techniques that allow for aptamer design and selection completely from scratch (**Gong *et al*., 2017**; **Hu *et al*., 2017)**. These include the computational design of an aptamer library **(Chushak *et al*., 2009; Lu *et al*., 2010)**, as well as a selection of aptamers against a given target **(Hu *et al*., 2015)**. The general problems with these approaches are the lack of random libraries and the necessary calculation power, which is disproportionately high **(Chushak *et al*., 2009; Hu *et al*., 2015; Prediger, 2016)**. Another interesting approach was published in 2016 by Ahirwar and colleagues who tried to design an aptamer to target estrogen receptor alpha (ERα) *in-silico*. Instead of generating a pool of random sequences, they deployed a “bottom-up” method utilizing a natural binding ligand of ERα— estrogen response element—to create aptamers and optimized the binding affinities **(Ahirwar *et al*., 2016)**. An interesting alternative approach was recently presented by a group from Heidelberg trying to circumvent the main flaws of aptamer design and development using experimental and *in silico* approaches. In 2015, the group participated in the International Genetically Engineered Machine (iGEM) competition and developed software named M.A.W.S., Making Aptamers Without SELEX. The algorithm loads the target molecule and creates a bounding box. For each nucleotide, the best conformation is calculated and the position with the lowest entropy is selected as the starting point for the next calculation cycle. Thereby, the algorithm constructs aptamers from scratch without using a given aptamer library against any given target structure **(Prediger, 2016)**.

Unlike other mentioned algorithms, the M.A.W.S algorithm was never tested for usability and applicability for aptamer generation for specific targets. Considering the low computational power needed for such algorithm to operate, we are proposing to attempt to use the M.A.W.S algorithm for an aflatoxin B1 specific aptamer generation.

## 4 Aflatoxins

Aflatoxins (AFs) are secondary metabolites produced by plant fungal pathogens infecting crops with strong carcinogenic and mutagenic properties. AFs are the predominant and most carcinogenic naturally occurring compounds which inevitably result in health complications, including hepatocellular carcinoma, acute intoxication, immune system disorder and growth retardation in children **(Groopman *et al*., 2008; Reddy *et al*., 2010)**. In 1993, AFs were classified as a Class I carcinogens by the International Agency for Research on Cancer (IARC) **(IARC, 1993; IARC, 2002)**. Among AFs, aflatoxin B1 (AFB1) is the most toxic and carcinogenic compound known. AFs are commonly relevant to the cereals, nuts and a scope of their agricultural products, especially peanuts, maize and rice **(Luttfullah and Hussein, 2011)**. In the United States, the Food and Drug Administration (FDA) has set the limiting value at 20 μg/kg for total AFs (B1, B2, G1, G2) for all foods, and 100 μg/kg for peanut and corn feed products. The European Commission (EC) set the maximum residue limit at 2 μg/kg for AFB1 and 4 μg/kg for total AFs **(Pan *et al*., 2020)**. The appearance of antifungal resistance of chemical fungicides and the safety requirements of practical application in crops globally have incurred the discovery of novel antifungal agents and other antifungal substances **(Bhatnagar-Mathur *et al*., 2015)**.

### 4.1. Producing microorganisms

Aflatoxins are produced as secondary metabolites by two fungal species, namely *Aspergillus flavus* (produce only B aflatoxins) and *Aspergillus parasiticus* (produce both B and G aflatoxins) **(Younis *et al*., 2020)**.

### 4.2. Toxicity

To date, at least 20 different types of AFs such as aflatoxin B1, B2, G1 and G2 have been identified. However, aflatoxin B1 (AFB1) is considered one of the utmost potent carcinogens, hepatotoxic, mutagenic, immunosuppressive, neoplastic, and teratogenic **(Asghar *et al*, 2014)**. Various environmental elements such as agricultural practices, moisture, temperature, unseasonal rains, geographical location, storage, and transportation are also favored the growth and production of *Aspergillus* and AFB1 contamination. The massive economic effects on the agriculture segment can also be considered because this contamination is able to decrease nutritional significance of food and feedstuff, reduction in meat production, reduced kidney and liver function, immune system suppression and finally created harmfulness to the consumers of dairy/ foodstuffs **(Asghar *et al*, 2020)**.

The mechanisms of AFB1-induced hepatotoxicity in S phase-arrested L02 cells using single-cell RNA-seq and single-cell reduced representation bisulfite sequencing (RRBS) was investigated and it was revealed that AFB1 induced apoptosis and cell cycle S phase arrest, reduced mitochondrial membrane potential (ΔΨm), and increased reactive oxygen species (ROS) generation, as well as the DNA methylation level. DNA methylation played a role via regulating the gonadotropin-releasing hormone receptor pathway, the Wnt signaling pathway, and the TGF-beta signaling pathway **(Zhang *et al*., 2020)**.

Aflatoxin B1 forms bulky DNA adducts and result in mutational spectra with typical C>A mutations, respectively, similar to those observed in exposed human cancers and cell lines **(Volkova *et al*., 2020)**. AFB1 also reduces intracellular ATP levels **(Zhu *et al*., 2020)**. Furthermore, AFB1 induced reduction in growth, feed utilization, digestive enzymes activities, and antioxidant enzyme activities; elevated the activities of liver enzyme, DNA damage, residual aflatoxin in Nile tilapia musculature **(Hassan *et al*., 2020)**.

ROS levels in IMR-32 cells increased significantly in a time-and AFB1 concentration-dependent manner, which was associated with the upregulation of NOX2, and downregulation of OXR1, SOD1, and SOD2. Substantial DNA damage associated with the downregulation of PARP1, BRCA2, and RAD51 was also observed. Furthermore, AFB1 significantly induced S-phase arrest, which is associated with the upregulation of CDKN1A, CDKN2C, and CDKN2D. Finally, AFB1 induced apoptosis involving CASP3 and BAX. Taken together, AFB1 manifests a wide range of cytotoxicity on neuronal cells including ROS accumulation, DNA damage, S-phase arrest, and apoptosis—all of which are key factors for understanding the neurotoxicology of AFB1 **(Huang, 2020)**. For these toxic effects, the European Commission set the maximum residue limits of aflatoxin B1 for various types of susceptible food summarized in **(Table 2.1.)**.

**Table 2.1.** Maximum Residue Limits in different foodstuffs according to **(EC, 2020)**.

| FOODSTUFFS | MAXIMUM RESIDUE LIMIT (MG/KG) |
| --- | --- |
| Groundnuts (peanuts) and other oilseeds | 8.0 |
| Almonds, pistachios, and apricot kernels | 12.0 |
| Hazelnuts and Brazil nuts | 8.0 |
| Tree nuts, other than the tree nuts listed | 5.0 |
| Groundnuts (peanuts) and other oilseeds for direct human consumption | 2.0 |
| Almonds, pistachios, and apricot kernels for direct human consumption | 8.0 |
| Hazelnuts and Brazil nuts for direct human consumption | 5.0 |
| Tree nuts, other than the tree nuts listed for direct human consumption | 2.0 |
| Dried figs | 6.0 |
| Dried fruit except figs | 5.0 |
| Maize and rice | 5.0 |
| All cereals | 2.0 |

### 4.3. Extraction

An amount of 0.2 to 63 mg of aflatoxins B1 and G1 per 100 ml from isolates of *Aspergillus flavus* in a nutrient solution consisting of 20% sucrose and 2% yeast extract was produced by (**Davis *et al*., 1966)**.

A method for the isolation of highly purified aflatoxins B from extracts of *Aspergillus flavus*. The aflatoxin, isolated from background impurities by rapid passage of the extracts through an acid alumina column, are separated from each other by chromatography on a silica get column, then separated by elution from a silica gel column with chloroform/Ethanol buffer was described by **(Rodricks, (1969)**.

Liquid–liquid instrumental separation, isolated the four aflatoxins, namely B1 (400 mg), B2 (34 mg), G1 (817 mg) and G2 (100 mg) successfully with 96.3%–98.2% purity from 4.5 L of *Aspergillus parasiticus* fermented material in a 250 mL centrifugal partition chromatography column was used by **(Endre *et al*., (2019)**.

Due to the apparent significance of detecting mycotoxins, specifically aflatoxin B1, designing an aptamer with high affinity for such disastrous toxin is of an utter importance.

## III. Materials and Methods

### 1. Materials

#### 1.1 Computational Aptamer design and generation

MAWS algorithm was used to generate high affinity DNA aptamer sequence against Aflatoxin B1. All computational methods were performed in an Anaconda Python environment, according to the maws.readme file of the M.A.W.S algorithm. Ambertools21 was used to generate PRMTOP files for the 4 different DNA nucleotides to be used in the MAWS 2015 algorithm **(Case *et al*., 2021)**.

#### 1.2. Aptamer enhancement through molecular docking simulations

A pre-existing aflatoxin B1 aptamer sequence with a dissociation constant of less than 2×10^−6^M (5′-TGG GCA CGT GTT GTC TCT CTG TGT CTC GTG CCC T-3′) **(Linda *et al*., 2011)**. The sequence will be modelled using the combination of Mfold web server, RNAComposer, Web 3DNA 2.0, and YASARA tools **(Krieger *et al*., 2009; Krieger and Vriend, 2014; Li, Olson and Lu, 2019; Popenda *et al*., 2012; Zuker, 2013)**. AutoDock Vina was used as the molecular docking tool and PyMol was used to analyze and visualize the results **(Trott and Olson, 2010; Schrödinger LLC, 2015)**.

### 2. Methods

#### 2.1. Computational aptamer generation for Aflatoxin B1

The M.A.W.S algorithm was downloaded from GitHub. The algorithm was run on a Linux-based system and windows to ensure convenience and applicability of the algorithm on different OS environments. The aflatoxin B1 small organic molecule was preprocessed by using antechamber algorithm to obtain a mol2 file. The sdf file of the molecule was obtained through the PubChem database. After the Aptamer is generated, it was synthesized by local DNA synthesizer.

The M.A.W.S algorithms, both 2015 and 2017 versions, were downloaded from GitHub. The algorithm was natively directed to work on a Linux-based system, however with minor adjustment it was also capable of running on Windows. The SDF and PDB files of the aflatoxin B1 molecule were obtained through the PubChem and EMBL-EBI databases, respectively. The aflatoxin B1 was preprocessed by using antechamber algorithm to obtain a mol2 file.

Crystal structures (PDB) files of the 4 DNA nucleotides were obtained through the EMBL-EBI database and Two files, PRMTOP and INPCRD, were obtained for every nucleotide using Ambertools21 **(Case *et al*., 2021)**. Mol2 files that are required by the LEaP tool used were created using these PDB files using the antechamber tool **(Case *et al*., 2021)**. To obtain all the parameters that were missing in the mol2 file, we used the parmchk tool and a frcmod file was obtained **(Case *et al*., 2021)**. The mol2 files and the amber parameters (frcmod files) were loaded in tleap.

#### 2.2. Enhancement of a pre-existing Aptamer computationally

Molecular modelling and docking were used, and according to analysis of the molecular docking simulations results, one point mutation was induced that influenced the aptamer 3D structure significantly. The ssDNA aptamer was modelled as follows, the secondary structure was predicted using Mfold web server for nucleic acid folding and hybridization prediction. RNAComposer was used to predict the 3D structure of the ssDNA after it was converted to RNA. To convert the resulted PDB file back to DNA, Web 3DNA 2.0 was used to mutate the uracil nucleotides back to thymine and YASARA was used to visualize and add hydrogens, and to minimize its energy. Molecular docking of the aflatoxin B1 ligand to the aptamer was performed using AutoDock Vina. The ligand and the aptamer were prepared using AutoDock tools and a two grid boxes were obtained for the aptamer and new mutated aptamer. The first grid box encompassed the entirety of the aptamer and the other one was directed at the most prominent binding site. Multiple simulations were performed to compare between the aptamer to the new mutated aptamer using the two gird boxes. The results were analyzed and visualized through PyMol.

## IV. Results

The soul aim of this work was the development of a novel aptamer that is has high affinity and specificity for aflatoxin B1 without to resort to carrying out the SELEX method. Our initial approach was to use the MAWS algorithm that was specifically written for that purpose, developing aptamers against proteins or small molecules without SELEX. However, despite our intensive efforts to make the MAWS algorithm operatable, it did not output any result whatsoever. A new approach then was needed to achieve the main goal. The new approach was comprised by molecular modelling of a preexisting aptamer, molecular docking, and analysis of that aptamer computationally and introducing point mutation to enhance that aptamer and then confirm that enhancement through comparing molecular docking results and through in vitro analysis.

### 1. Operating the MAWS algorithm

It has been claimed that the MAWS program will be ran smoothly after following the instructions (readme) file. What was experienced was the exact opposite. Number of errors was raised during the use of the algorithm. The algorithm itself has two versions, MAWS 2015 and 2017. MAWS 2017 was much less popular, hence the difficulty of obtaining it from GitHub as it required an intensive keywords search through several search engines in order to obtain the program itself. The main reason for that difficulty is that the creators uploaded the file and named it “sharksome-suite”, rather than MAWS and it can be found in this repository https://github.com/igemsoftware2017/AiGEM_TeamHeidelberg2017.

The newer version of the program was modularized and the dependencies of MAWS 2015 was removed. However, running MAWS 2017 was not successful at all, it raised many errors that we believe mainly originated from the fact that MAWS 2017 was specifically developed for proteins and not small molecules which was problematic because aflatoxin B1 is a small molecule and not a protein. MAWS 2015 did not have that issue as it has instructions on how to use it for proteins and small molecules. However, MAWS 2015 raised “file missing” error that prevented it from launching. The error was raised when the program tried to choose a nucleotide from the DNA nucleotides library randomly to begin the seed and grow process. The missing files where PRMTOP and INPCRD files of the 4 DNA, which are parameter/topology and coordinate files, respectively. Even though that error was resolved by creating the 8 missing files (2 files/nucleotide) using Ambertools21 as described previously, a new error emerged.

The new error was a “module not found” error, the module name was SciPy which is a source software for mathematics, science and engineering and is essential for scientific computing. The module was installed, and the error was resolved.

A new error was raised, an index error. The list of “best positions” of the nucleotides was completely empty, which only means that the software did not record any positions in it. The error here lies in the core code of the algorithm, in order to resolve such error, we tried to contact the programmers who wrote the algorithm. They were out of reach.

### 2. Molecular modelling and docking for aptamer enhancement

The ultimate aim of this work was to generate a new aflatoxin B1 aptamer computationally. Giving the errors that were generated by the MAWS algorithm, we decided to shift our focus into a new method. This new method depends on molecular modelling and docking to enhance a preexisting aptamer.

The DNA aflatoxin B1 aptamer, here named APB1-49, was modelled through a new pipeline that was not used in the literature prior to this work **(Figure 4.1)**. There is no direct way to model a single stranded DNA aptamers, the general concept is to model it as a single stranded RNA then convert the 3D structure obtained into a single stranded DNA 3D structure. To achieve that, we used Mfold, a web server for nucleic acid folding and hybridization prediction, to predict the secondary structure of the DNA. RNAComposer was used to predict the 3D structure of the ssDNA after it was converted to RNA. Mutation is needed to replace all Uracil nucleotides of the RNA back to Thymine. To achieve that Web 3DNA 2.0 was used. YASARA was used to visualize the PDB and add hydrogens, and to minimize its energy.

**Figure (4.1).**
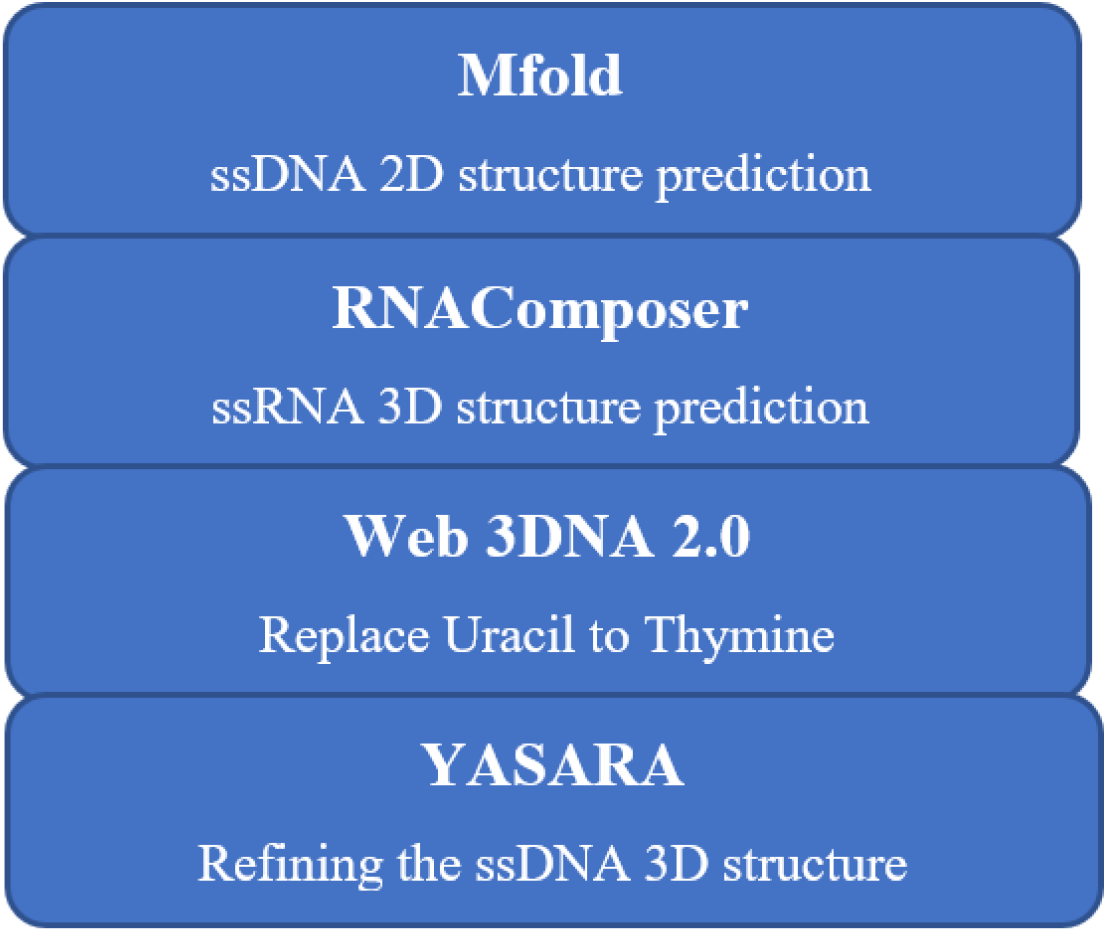
A new pipeline for ssDNA aptamers modelling.

The result was a PDB file that represents the aptamer 3D structure. The PDB file was then visualized by PyMol **(Figure 4.2)**. The PDB file was then used in AutoDock Tools in order to prepare it for the molecular docking process as a macromolecule. The result was a PDBQT alongside a grid box text file. The toxin was also prepared by AutoDock Tools as a ligand.

**Figure (4.2).**
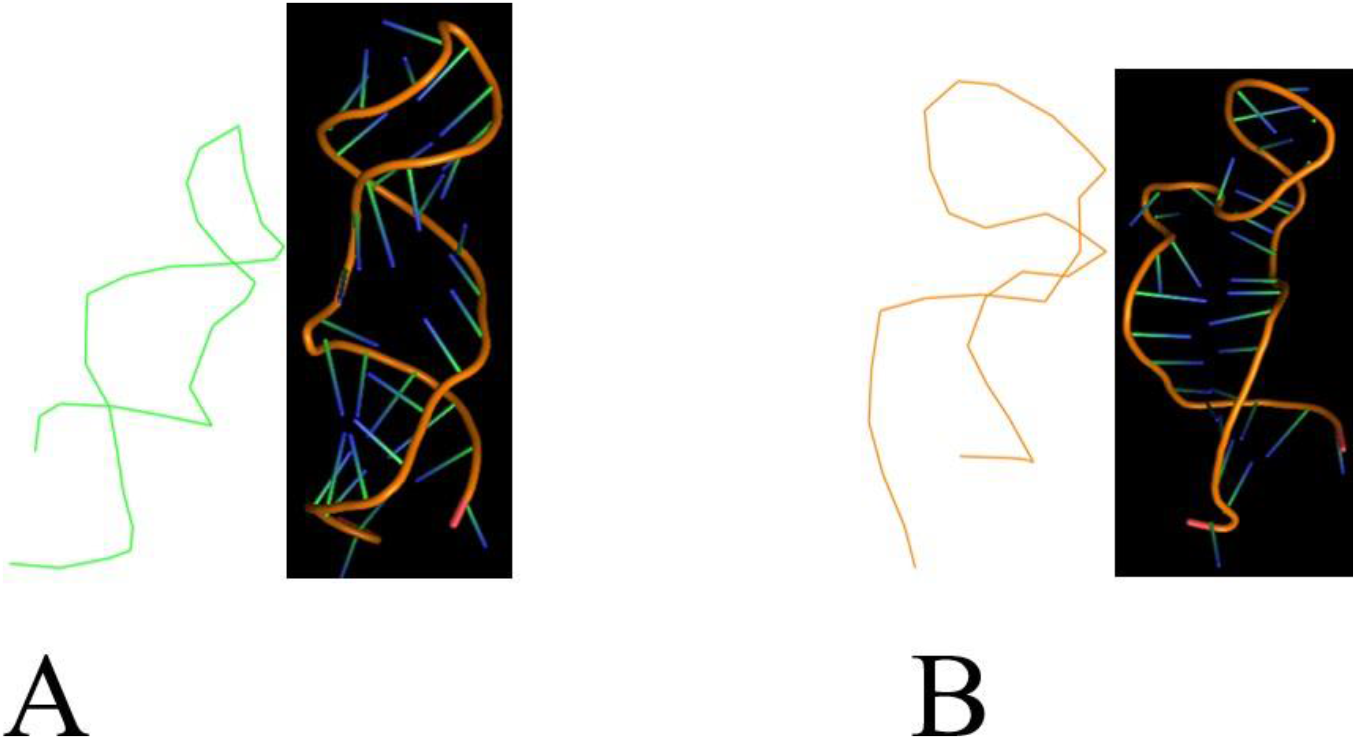
Comparison between 3D structures of the original and mutated aptamer. (A): 3D structure of the mutated aptamer, (B): 3D structure of the original aptamer.

Four runs of docking sessions were performed, and the results were recorded and positions that have RSMD value bigger than 10 were excluded. The grid box that was used covered all of the APB1-49 molecule and then the resulted positions were visualized using PyMol to further downsize the grid box into a specific pocket area. The process was repeated using the new grid box coordinates and results that have RSMD value bigger than 10 were further excluded.

The analysis of hydrogen bonds and Van Der Waals forces of the positions obtained allowed us to introduce one point mutation that showed potential enhancement of the current aptamer affinity to the toxin. Accordingly, we introduced one point mutation of 28^th^ nucleotide (G → C), using Web 3DNA 2.0. The mutated aptamer, APB1-49M, was modelled and molecular docking was performed as previously discussed. The final results were grouped, inspected, and analyzed **(Figure 4.3)**. The hydrogen bonding between the ligand and the original aptamer was analyzed and compared with the mutated aptamer hydrogen bonding with the ligand **(Figure 4.4)**.

**Figure (4.3).**
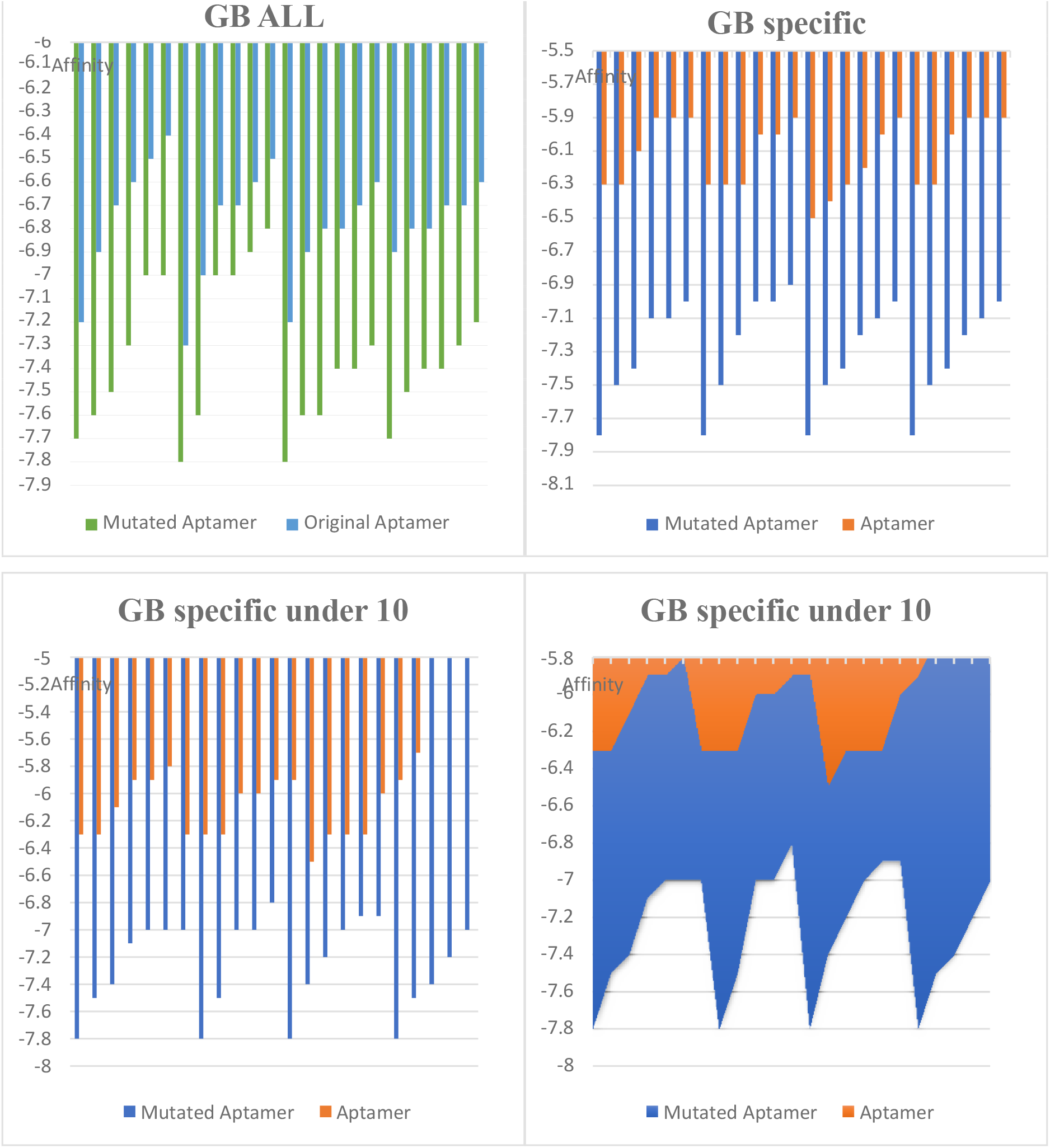
Comparison between the average of Vina docking scores between the mutated aptamer and the original one. (GB ALL) → Grid box covered all the aptamers, (GB specific) → Grid box covered the same most plausible pocket in the aptamers, (Under 10) → Only scores with RSMD of less than 10 was considered.

**Figure (4.4).**
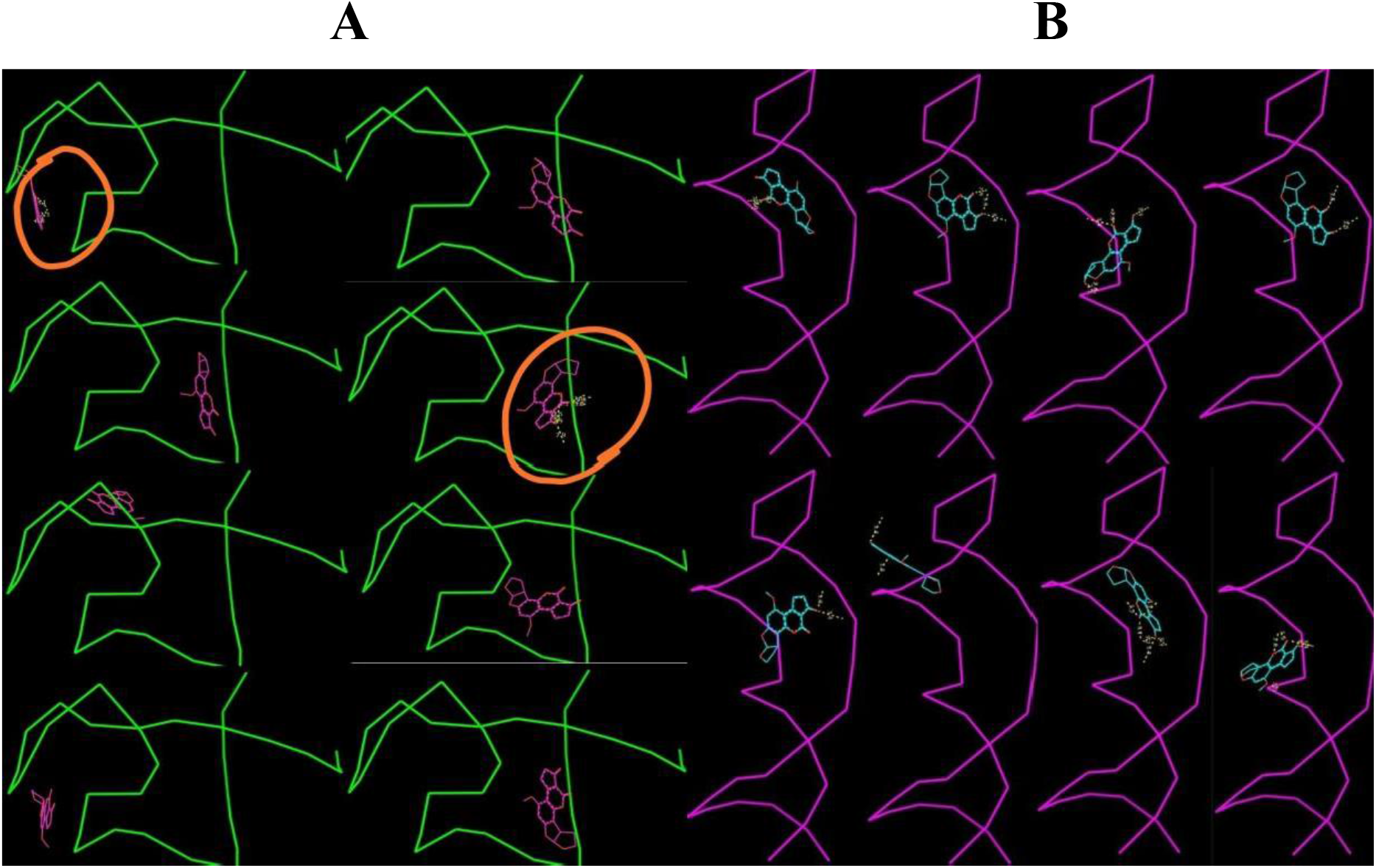
Comparison between the existence of hydrogen bonds between the ligand (Aflatoxin B1) and the original aptamer (Green) and between the mutated aptamer (Purple). (A) Only two poses showed hydrogen bonding between the ligand and the original aptamer (Circled). (B) Shows hydrogen bonding between the ligand and the mutated aptamer in all of the 8 poses.

## V. Discussion

Despite being mentioned in the literature and our vigorous efforts, the MAWS algorithm did not function as was claimed by its creators. The principle behind the MAWS algorithm, the seed and grow method, is valid and we would recommend adopting such principle in order to develop a functioning version of the MAWS algorithm for designing *de novo* aptamers for proteins and small molecules ligands. However, until a functioning version of MAWS is developed, our focus was directed to the usage of molecular modelling and molecular docking to develop enhance an existing aptamer against Aflatoxin B1.

The modelling of ssDNA aptamers is quite difficult. Current existing tools are for the modelling of double stranded DNA or single stranded RNA. Hence the need for an extremely easy and fast pipeline to model single stranded DNA aptamers. The general principle is predicting the secondary structure of the DNA, then treat the ssDNA as ssRNA in order to predict its 3D structure. After obtaining the 3D structure in the form of PDB file, the next step should be changing it from RNA back to DNA. Several pipelines were proposed in the literature, however, most of them rely heavily on manually changing all uracil nucleotides back to thymine nucleotides, which requires a lot of effort, then minimize the energy of the 3D structure to adequately suit the nucleotide change.

Here, we provided a new pipeline that is uniquely using a new tool, web 3DNA 2.0, that automatically and easily replace all uracil nucleotides back to thymine nucleotides **(Figure 4.1).)**. Moreover, all the tools were used are web servers, which means that there will be no need for any kind of installation. The time required for the modelling of one ssDNA aptamer was about 5 minutes to 30 minutes, the time required of every web server did not exceed 30 seconds using adequate internet connection, except for the YASARA energy minimization server. The YASARA server required 4 to 30 minutes, in order to obtain the minimized PDB file via email. One can circumvent such delay by installing their program, which will result in obtaining the minimized PDB file in less than a minute, depending on computational power of the used machine.

The visualization of the docking results by PyMol was of a great worth in determining the point mutation needed to significantly enhance the affinity of the aptamer against the ligand. This further proves that the use of molecular docking is a great viable option in aptamer design and development.

The analysis of the results, of both the docking scoring and hydrogen bonding between the molecules, has led us to the determination that APB1-49M is a greatly enhanced version of APB1-49. This is for two main reasons, the first one is that the affinity score of AutoDock Vina suggested that the mutated aptamer APB1-49M has more affinity against aflatoxin B1 than the original aptamer APB1-49 **(Figure 4.3)**. The second reason is that only two of eight docking poses had hydrogen bonding between APB1-49 and AFB1, while eight of eight poses had hydrogen bonding between the APB1-49M and AFB1 **(Figure 4.4)**. Further analysis of these hydrogen bonds showed that the length of the hydrogen bonds between APB1-49M and AFB1 was shorter than that between APB1-49 and AFB1, which means it is more stable.

This suggests that APB1-49 primarily depend on Van der Waals forces to bind with AFB1, while APB1-49M depends on the stronger type of bonding, hydrogen bonds. Even though these *in silico* results are in favor of that determination, there is a need to confirm these results *in vitro*.

## VI. Conclusion

The presence of mycotoxins, especially Aflatoxin B1, in food and feed is very dangerous and therefore the detection of such toxins is of the upmost importance. Biosensors are being used to detect these toxins quickly and easily, which saves effort, money and time. Aptamers are an excellent choice to be used as biorecognition elements as they hold high affinity and specificity for target molecules, can be easily manipulated and modified, more stable than antibodies (especially DNA aptamers), does not need immune reactions to be generated, and they are much cheaper to be obtained. SELEX remains the main method to obtain aptamers against ligands of interest, however it is a laborious process that consumes an unnecessary amount of time and effort. Various efforts were done during the past two decades to obtain aptamers computationally without performing SELEX cycles.

The MAWS program was one of these efforts, and the principle behind it, the seed and grow method, is outstanding. However, the program failed to function properly. The creation of an algorithm based on the same basic principle but that functions appropriately and updated regularly is greatly recommended.

Despite the MAWS program failure, we were able to generate a new aptamer against Aflatoxin B1 computationally without resorting to SELEX, using only molecular modelling and molecular docking. This new aptamer is an enhanced version of a preexisting aptamer.

Tools for ssDNA aptamers 3D structure prediction are lacking. Therefore, a new pipeline was proposed to accurately predict the 3D structure of ssDNA aptamers, that is easier and more effortless than current proposed pipelines in the literature. We recommend the implementation of this pipeline into one algorithm for aptamers (DNA or RNA) 3D structure prediction.

The molecular docking simulations were performed using AutoDock Vina and the results were visualized using PyMol. The examination of the docking results is in favor of that claim, and we are recommending to further prove it using *in vitro* analysis of the APB1-49M aptamer affinity and dissociation constant and compare it with the APB1-49 aptamer.

## Supporting information

Acknowledgments and Supp data

