## Supplementary material for "Computational Design of a New Aflatoxin B1 Aptamer *in lieu* of SELEX Technique": Acknowledgments and Supp data

### List of Figures


| Figure | Title | Page |
|---|---|---|
| **Figure 1.1** | Biosensors vs conventional techniques for mycotoxins analysis and detection. | **2** |
| **Figure 1.2** | Simple representative diagram of Biosensors general composition | **3** |
| **Figure 4.1** | A new pipeline for ssDNA aptamers modelling. | **35** |
| **Figure 4.2** | Comparison between 3D structures of the original and mutated aptamer. | **35** |
| **Figure 4.3** | Comparison between the average of Vina docking scores between the mutated aptamer and the original one. | **36** |
| **Figure 4.4** | Comparison between the existence of hydrogen bonds between the ligand (Aflatoxin B1) with the original aptamer and with the mutated aptamer. | **37** |

### List of Tables


| Table | Title | Page |
|---|---|---|
| **Table 2.1** | Maximum Residue Limits in different foodstuffs according to. | **19** |

### List of Abbreviations

| Abbreviation | Meaning |
|---|---|
| AF | Aflatoxin |
| AFB1 | Aflatoxin B1 |
| BWAs | Biowarfare agents |
| ERα | Receptor alpha |
| FACS | fluorescence-activated cell sorting |
| FAO | Food and Agriculture Organization |
| FFT | Fast Fourier Transform |
| FLD | Fluorescence detector |
| FT-NIR | Fourier Transform near-infrared |
| GC-MS | Gas Chromatography- mass spectrometry |
| IARC | International Agency for Research on Cancer |
| LC-MS/MS | Liquid Chromatography- Mass spectrometry |
| LOD | Limit of detection |
| M.A.W.S. | Making Aptamers Without SELEX |
| MMC | Mutation Monte Carlo |
| NMR | Nuclear magnetic resonance |
| PPB | Part per Billion |
| RRBS | Reduced representative bisulfate sequencing |
| SC | Shape complementarity |
| SDM | Sulfadimethoxine |
| SELEX | Systemic evolution |
| SPR | Surface plasmon resonance |
| VEGF | vascular endothelial growth factor |
| ZEN | Zearalenone |

### Acknowledgements

I wish to express my infinite indebtedness to Assist. Prof. Dr. Moez Elsaadani, Assistant Professor at faculty of Biotechnology, Misr University for Science and Technology, for his continuous support and his meticulous supervision. His suggestions have been of great help. His kind brotherhood has always been overwhelming.

I am endlessly grateful to Prof. Dr. Ahmed Z. Abdel Azeiz, Professor of Biochemistry, Faculty of Biotechnology, Misr University for Science and Technology, I will always be impressed with his knowledge, wisdom, and farsightedness.

I have the deepest gratitude for Marina Maurice Michel, Assistant lecturer at faculty of Biotechnology, Misr University for Science and Technology, she was there at every step of the way, helping me whenever her help was needed. Her faithful support made this work possible.
